# To be a Grid Cell: Shuffling procedures for determining “Gridness”

**DOI:** 10.1101/230250

**Authors:** C. Barry, N. Burgess

## Abstract

Grid cells in freely behaving mammals are defined by the strikingly regular periodic spatial distribution of their firing. The standard method of identification calculates the “Gridness” of the spatial firing pattern, with significance being defined relative to the 95^th^ percentile of a null distribution of the Gridness values found after randomly permuting spike times relative to behaviour. We determined the false-positive rate by applying the method to simulated firing with irregular spatially inhomogeneity (i.e. randomly distributed Gaussian patches). We found surprisingly high false positive rates (potentially approaching 20%), which were strongly dependent on the type of Gridness measure used and the number of spatial fields in the synthetic data. This likely reflects the spatial homogeneity of the distributions of spikes after shuffling compared to the inhomogeneous synthetic data. However, false positive rates were reduced (generally below 8%), and less dependent on other factors, when an alternative spatial field shuffling method was used to generate the null Gridness distribution. For comparison, we analysed single unit recordings made using tetrodes implanted into rat medial entorhinal cortex for the purpose of finding grid cells. We found 24% of active neurons were classified as grid cells via spike shuffling and 22% via field shuffling. These results, and the potentially high false-positive rate when classifying cells with patchy but irregular firing as grid cells, indicate that the proportion of cells with regular grid-like firing patterns can be over-estimated by standard methods.

## Introduction

Grid cells have been identified in the medial Entorhinal Cortex (mEC) of a variety of mammalian species^1–5^. Recorded in behaving animals, they show a spatial distribution of firing as a function of location that is characterized by a remarkable regular periodic structure. Here we examine the standard methods for identifying the spatially regular firing patterns of grid cells, focussing on the false positive rate when applied to neural firing that may include irregular spatial inhomogeneity, as is often seen in recording from animals whose behaviour is inhomogeneously distributed in space.

Grid cells were discovered in rat mEC ^4,6^ and, in the open field, fire whenever the animal enters an array of locations arranged across the environment at the vertices of a regular triangular grid. The standard method of identifying grid cells is based on a “Gridness” measure which quantifies the 6-fold rotational symmetry in the firing pattern ^7^. The spatial autocorrelogram (SAC) is calculated from the firing rate map, the annulus containing the six peaks nearest the origin is found and the firing rates in the annulus are correlated with those in rotated versions of the annulus. Grid cells should have a greater correlation for 60° and 120° rotations than for 30° 90° and 150° rotations. The “standard Gridness” measure is thus defined as the minimal (60°, 120°) correlation minus the maximum (30°, 90°, 150°) correlation ^7^. Subsequent modifications include “expanding Gridness” in which the radius of the annulus is varied and the maximum Gridness score is used^8^, “elliptical Gridness” in which the annulus is allowed to be elliptical^9^ and “expanding elliptical Gridness”^3,10^. The significance of a neuron’s Gridness is judged relative to the distribution of Gridness scores in a “spike-shuffled” population of firing rate maps used to simulate the null hypothesis of no grid cells. By rotating the times of the spikes relative to the locations ^8,11^ populations of firing rate maps are formed in which there is no spatial structure while maintaining temporal structure (Figure 1).

**Figure 1.**
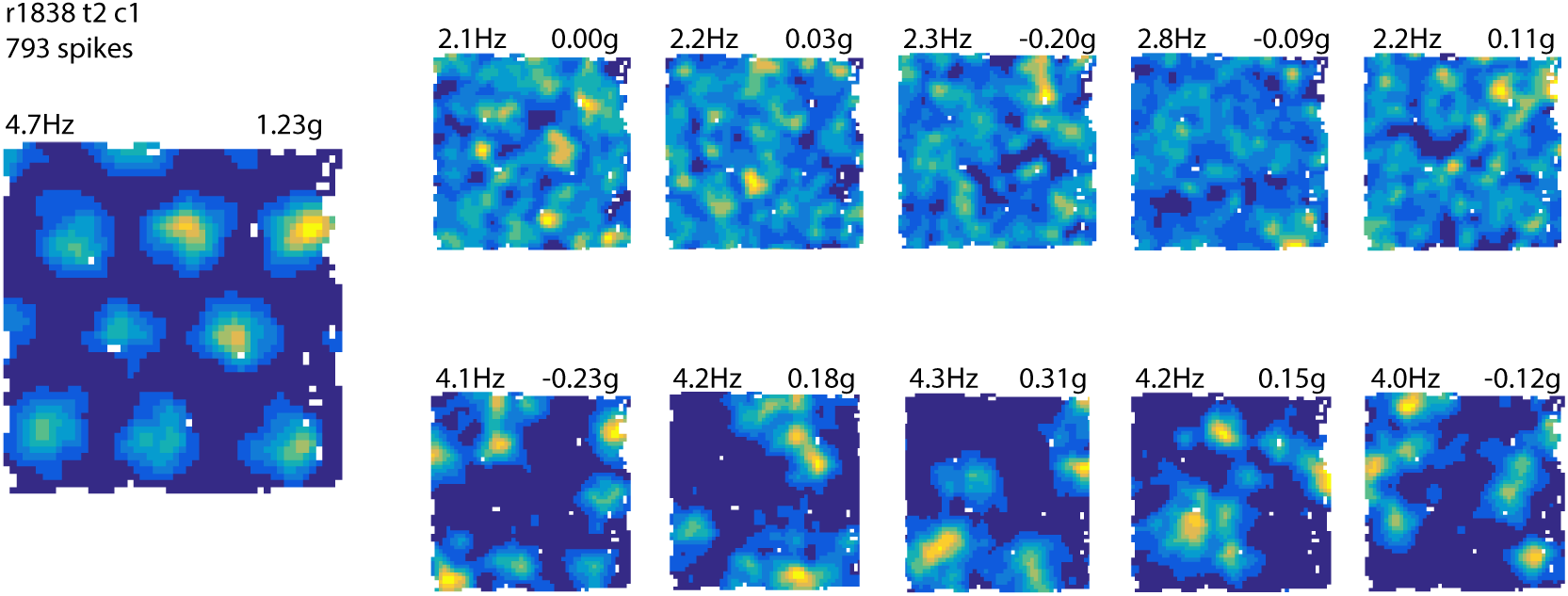
Effect of spike and field-shuffles on spatial firing. Left, firing rate map for a mEC grid cell. Peak rate and standard Gridness score are shown above the ratemap. Right top, five firing rate maps derived from spike-shuffled permutations of the grid cell firing. Firing maps are relatively homogeneous and exhibit low Gridness scores. Bottom, ratemaps derived from a field shuffling procedure. The local topography of fields is preserved while their global topography is perturbed. The range of firing rates and the inhomogeneity of their spatial distribution better matches the original map.

We were concerned that these methods might overestimate “Gridness”, because “spike-shuffled” data tends to be more uniformly distributed in space than is typical of data from freely behaving animals, in which the distributions of behaviour and sensory stimuli create inhomogeneity. One potential solution is to perform “field shuffling”, that is, allow inhomogeneity by having local clusters of firing, but randomise their locations relative to each other^12^. To investigate these issues, we analysed artificial datasets constructed to contain irregular spatial inhomogeneity - randomly distributed Gaussian patches of firing - so as to be able to estimate the false-positive rate for detecting grid cells in these circumstances. We then analysed datasets recorded from mEC in foraging rats^13,14^ to estimate the actual proportion of neurons with regular grid-like firing patterns that are detected by the various methods.

## Materials and Methods

### Neural data

#### Animals and surgery

Seven male Lister Hooded rats (250-402g at implantation) each received a single microdrive carrying four tetrodes of twisted 17-25 μm HM-L coated platinum-iridium wire (90% - 10%) (California Fine Wire, USA) targeted to the right dorsal medial entorhinal cortex (mEC). 17 μm wire was platinum plated to reduced impedance to 200-300kΩ at 1kHz. Surgical procedure and housing conditions were the same as those described previously by Barry et al^13^. Data has been previously analysed in^13,14^ All experiments were carried out in accordance with the U.K. Animals (Scientific Procedures) Act 1986.

#### Recording

Training and screening was performed post-surgically after a one week recovery period. An Axona recording system (Axona Ltd., St. Albans, UK) was used to acquire the single unit and positional data. For details of the recording protocol see^13,14^. The position of the rat was captured using an overhead video camera to record the position of the one or two LEDs on the animal’s head-stage. The animal’s head direction was extracted using the relative position of the two LEDs, one large, one small, positioned 8cm apart at a known angle to the rat’s head.

All animals were trained to forage for sweetened rice in a 1m by 1m square environment which consisted of a clear Perspex floor and 50cm high Perspex walls fronted with grey card (north and south walls) and black ribbed card (east and west walls). Training consisted of at least five trials each of 20 minutes, distributed over three days. Between trials the floor of the arena was wiped with a damp cloth to remove faeces, urine and uneaten rice. All subsequent recordings took place in this same familiar arena.

Multiple recordings were made across days with the electrodes advanced by 50μm between recordings. Each recording being 20 minutes during which animals foraged for rice. In situations where electrodes were not advanced, the first recording made with the electrodes in a specific configuration were submitted to further analysis. Recordings were typically ceased after grid cells were no longer detected.

#### Histology

At the end of the experiment rats received an overdose of Euthatal (Sodium pentobarbital) and were transcardially perfused first with phosphate buffered saline and then with 4% paraformaldehyde (PFA) solution. The brains were removed and stored in 4% PFA for at least one week prior to sectioning. 40 μm frozen sagittal sections were cut, mounted on gelatine-coated glass slides and stained with cresyl violet. The depth and layer at which cells were acquired was extrapolated by reference to the record of tetrode movements after taking account of brain shrinkage. All animals were confirmed to have successful recordings made from mEC.

### Data analysis

Spike sorting was performed offline using a data analysis suite (Tint, Axona Ltd., St. Albans, UK) and further analysis was conducted using Matlab (Mathworks, Natick, Mass. USA). Action potentials were assigned to putative cells based on amplitude, waveform and temporal autocorrelation criteria applied elsewhere to entorhinal grid cells^4,13^. In each recording session all unique cells, including those without obvious spatial correlates, were cut. Since electrodes were typically implanted above the mEC, only cells recorded after that first grid cell was identified were submitted to further analysis (n=704).

Position and concomitant spikes were binned into 2 × 2cm bins. Smoothed firing rate maps (‘ratemaps’) were calculated by dividing the number of spikes assigned to a bin by the cumulative occupancy of the bin after first convolving both the spike and dwell time maps with a Gaussian kernel (σ=1.5bins).

#### Standard Gridness

Standard Gridness was computed by first generating the spatial autocorrelogram of the ratemap ^7^. The autocorrelogram is defined by Pearson’s product moment correlation coefficient with corrections for edge effects and unvisited locations. With *λ*(*x*, *y*) denoting the average rate of firing of a cell at location(*x*,*y*), the autocorrelation between the fields with spatial shifts of τ*_x_*and τ*_y_* was thus estimated as:

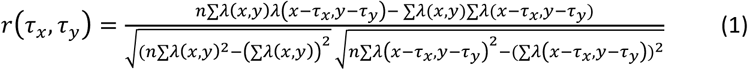

Then from the resulting spatial autocorrelogram we found the central peak and its surrounding field - defined as the region bounded by the peak’s half height - as well as the six closest peaks and fields. These peaks were used to define an annulus with the inner edge being the boundary of the central peak and the outer edge being a circle with the minimum radius that would fully encompass the fields of the six closest peaks. The region of the autocorrelogram bounded by this annulus was then rotated by increments of 30° up to 150° and for each rotation a Pearson product moment correlation coefficient was computed with the un-rotated mask. Gridness was then defined as the difference between the lowest correlation with rotations of 60°and 120° and the highest correlations of rotations with rotations of 30°, 90° and 150°.

#### Expanding Gridness

Expanding Gridness^8^ was computed by defining multiple circular annuli from the autocorrelogram, with outer radii increasing in steps of 1 bin (2.0cm) from a minimum of 10 cm more than the radius of the central peak, to a maximum of 10 cm less than the width of the environment. Gridness was then defined, as before, for each annulus and expanding Gridness was taken as the maximum score obtained for each cell.

#### Elliptical Gridness

Elliptical Gridness took the same form as standard Gridness following an ellipse fitting and regularisation step^9^. This measure compensates for single axis distortions in the grid-pattern by fitting an ellipse through the six central peaks of the autocorrelogram, finding the transform required to regularise that ellipse, and finally applying that transform to the entire autocorrelogram before calculating Gridness as before. First a linear least squares procedure is applied. The conic equation of an ellipse is considered:

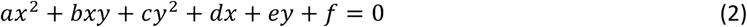

Then constructing a matrix from at least 5 points where each column takes the form:

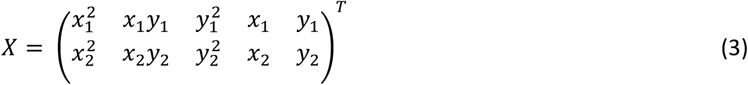

And the parameters vector 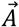 is constructed as:

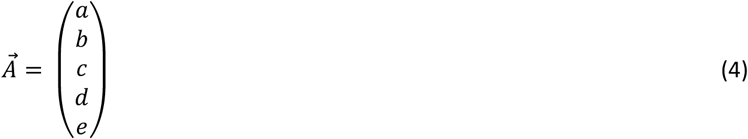

With the target vector 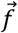:

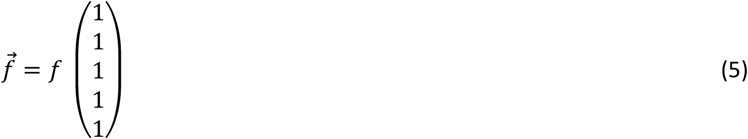

Therefore minimizing 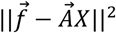 with respect to 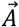 gives the standard linear least squares solution:

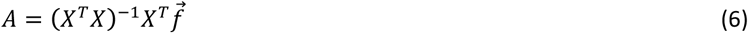

Once we estimate these parameters it is trivial to calculate the orientation and length of the minor and major axis of the ellipse and we use this information to resize the autocorrelogram such that the major axis is compressed to the length of the minor axis.

Following this regularization, standard Gridness was calculated on the transformed autocorrelogram. In situations where fewer than six peaks were found to surround the origin, the regularisation step was not applied and standard Gridness was calculated in lieu.

#### Expanding elliptical Gridness

Expanding elliptical Gridness was generated by first applying the regularisation step described for elliptical Gridness followed by the procedure for calculating expanding Gridness ^3,1^.

#### Shuffling Procedures

Two permutation methods were applied in order to generate null distributions against which the significance of Gridness scores were tested: spike shuffling and field shuffling. In each case data from each cell was used to establish a threshold for that cell. Specifically, for each cell, the shuffling procedures were applied 100 times, generating a population of ratemaps. The Gridness scores of these ratemaps were calculated, forming a null distribution. Finally, the 95^th^ percentile of these distributions were used to set a significance threshold, such that cells with Gridness exceeding this threshold were classified as grid cells.

##### Spike shuffling

Spike shuffling was performed as described in ^8,11^. Specifically the spike train for each cell was permuted in time by a random number ∊ [20, *d* — 20] seconds where *d* is the trial duration. The new time indices of spikes which exceed the trial duration were wrapped around to the beginning of the spike train. The effect of this procedure was to preserve the temporal characteristics of the spike train – the inter spike intervals - while disrupting the relationship between spikes and an animal’s spatial behaviour (which was indexed by the spike time).

##### Field shuffling

Spatial shuffling^12^ provides a means to disrupt the long-range spatial topography of a ratemap while preserving, as far as possible, the short-range topography of the fields. First, fields within each ratemap were segmented by applying a watershedding procedure to a highly smoothed version of the ratemap (2cm bins, Gaussian kernel σ=3bins). The bin with the highest firing rate – the peak bin - within each field was identified. Subsequent steps were performed on ratemaps that had received a standard level of smoothing (Gaussian kernel σ=1.5bins). The peak bin of each field was copied to a random position within the corresponding shuffled ratemap. Bins around each peak were then copied to the shuffled ratemap, retaining as far as possible their proximity to their peak bin. This procedure was applied incrementally to each field as follows. The fields were numbered in a random order, then starting with the first field a bin immediately adjacent to the peak bin was selected from the original ratemap and placed, as far as possible, in the same relative position to the peak bin in the shuffled ratemap. If this location was not available or already occupied the next nearest bin was used. The same procedure was then applied to the closest bin to the peak of each field. When a single bin had been replaced from each field, the second closest bins were replaced in the same order. Ultimately the procedure was repeated until all bins within the original ratemap had been copied to the shuffled ratemap. Unvisited bins were not moved.

### Simulated Inhomogeneous Spatial Firing

We created synthetic non-periodic spatial firing by randomly placing spatial fields – modelled as summed symmetric 2-d Gaussians - into a 1m square enclosure. The number and size of fields were chosen to correspond to grid cells with scales of 20 to 80cm, this range was explored in 5cm increments.

Specifically, the standard deviation of the Gaussians was defined as:

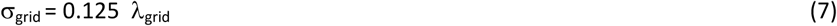

where *λ*_grid_ is the grid scale being matched, and the number of fields was defined as:

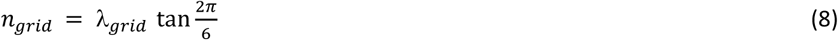

For each grid scale 1000 synthetic ratemaps were generated, with the peak firing rate being 10Hz. Next, to convert these idealised ratemaps to time series of neural data with matching behaviour, we used positional data derived from ten 20 minute trials in which mEC implanted rats foraged for rice in a familiar 1m square enclosure^14^. Positional data was sampled at 50Hz, thus each 20 minute trial yielded 60,000 (x,y) pairs, indicating the animals’ locations during the trials. These data were split into 1 minutes chunks, of 3,000 (x,y) pairs, and for each synthetic ratemap, 20 minutes of behaviour was generated by drawing at random twenty 1 minute chunks and concatenating them. To generate corresponding spike time series, the surrogate positional data was subsampled at 2ms intervals and at each interval the animal’s position used to determine an expected firing rate and thus expected number of spikes in the interval.

Finally, a Poisson point process was used to determine if 1 or 0 spikes was present in each interval. The synthetic data generated in this way was then binned into ratemaps and processed as described above in order to determine the Gridness of the simulated spatial fields. The significance of the Gridness measure was then assessed relative to spike and field shuffled null distributions derived from the same data. Thus, each simulated cell was categorised as a grid cell or not and the proportion of ‘grid cells’ of each scale were calculated.

## Results

### Distributions of Gridness in spike-shuffled and field-shuffled data

To illustrate the effects of spike shuffling and field shuffling we use a single unit from dorsomedial entorhinal cortex, which clearly exhibits a regular triangular symmetry (i.e. a grid cell). Figure 1 shows the effects of spike shuffling and field shuffling on the firing rate maps of this cell. Figure 2 shows the distributions of Gridness found in the populations of firing rate maps created by spike shuffling shuffling and field shuffling shuffling of the data from the two cells shown in Figure 1. The distributions of Gridness are shown for the four different measures of Gridness currently in use.

**Figure 2.**
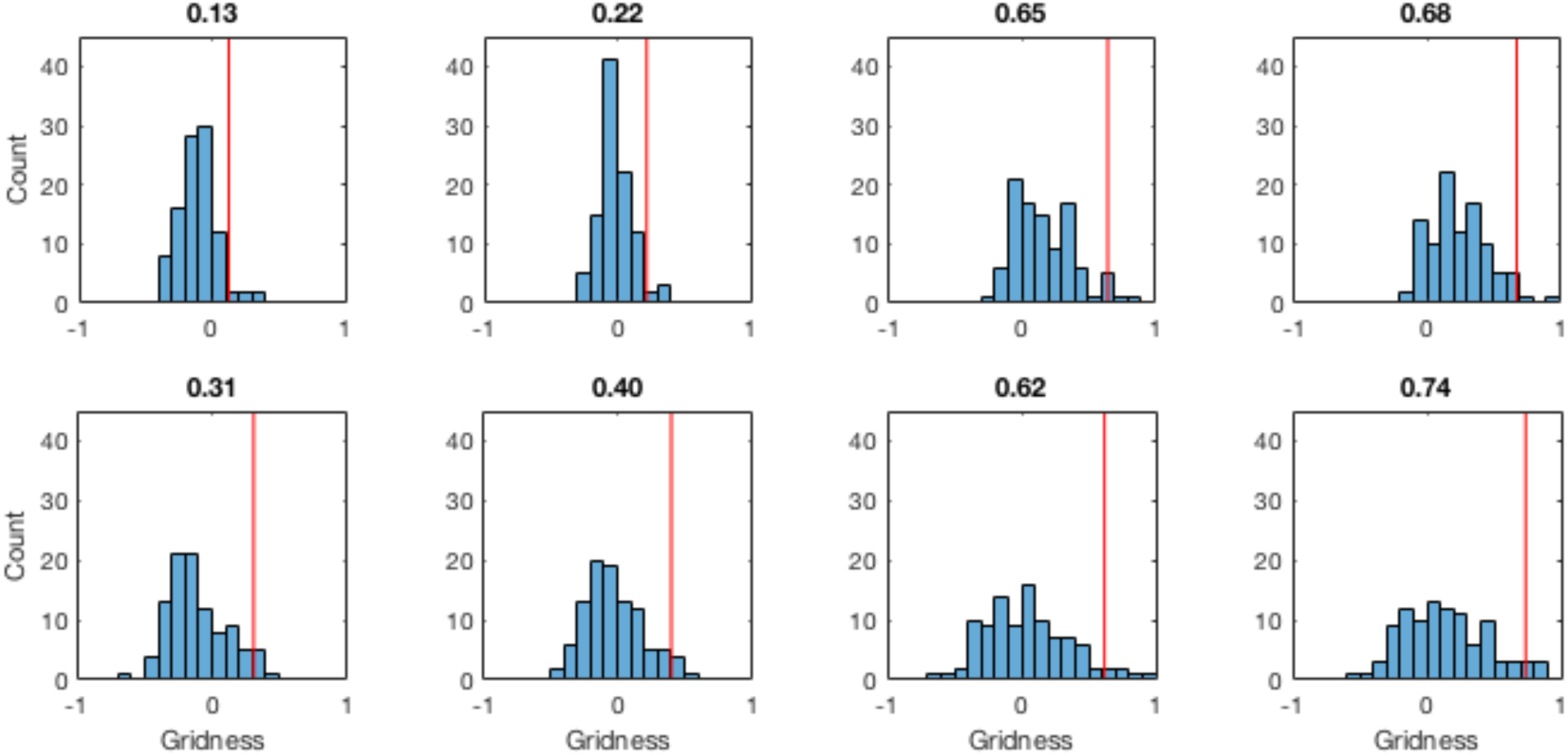
The distribution of the four measures of Gridness generated by spike shuffling (top row) and field shuffling (bottom row) of the data from the cell in Figure 1. Each distribution is generated from 100 shuffles of the data and the Gridness measures (columns) are in left to right order: standard Gridness^7^, elliptical Gridness^9^, expanding Gridness^8^, and the combination of expanding and elliptical Gridness ^3^. The title of each plot indicates the 95^th^ percentile threshold extracted from each distribution, typically used to identify grid cells.

Figure 2 illustrates our concern regarding the use of spike shuffled null distributions. The grid cell classifies as a grid cell relative to both spike shuffled and field shuffled distributions - its Gridness value exceeds the 95^th^ percentile of both shuffled distributions. However, the thresholds obtained by the different shuffling procedures varies considerably. For example, the standard Gridness threshold (Figure 2, column 1) obtained from a spike shuffle was 0.13 whereas a field shuffle yielded a threshold of 0.31 (Figure 2, top and bottom respectively). Clearly some cells that are identified as grid cells relative to the spike shuffled threshold would not exceed a threshold identified from a field shuffling procedure. Thus, there is likely to be a difference in the false positive rates obtained by these different approaches.

Similarly, the absolute values of Gridness vary considerably according to which version of the Gridness measure is used (Figure 2). Gridness values are higher for the expanding Gridness measure (column 3), as it takes the maximum Gridness value over several sizes of annulus, and for the elliptical Gridness measures (column 2) as any elliptical distortion in the data is “corrected” to circular before Gridness is calculated. The combination of both elliptical and expanding Gridness (column 4) yields the highest values. Thus, absolute values of Gridness cannot be compared between papers that use different versions of the Gridness measures, and “rules of thumb,” such as accepting regular Gridness scores exceeding fixed values will be much more lenient if used with expanding or elliptical Gridness, with expanding elliptical Gridness being the most permissive.

### The Gridness of simulated spatially irregular firing patterns

We simulated inhomogeneous irregular firing patterns to test the false-positive rate of grid cell identification. Spatially irregular Poisson firing patterns were generated from spatial firing rate distributions formed from summed randomly placed Gaussians (see Methods for details). Specifically, ratemaps corresponded to 1m × 1m square enclosures, with the size and number of fields within the enclosure chosen to match grid cells of scales between 20 and 80cm ^7^.

Figure 3 shows the proportions of simulated spatially irregular cells that are classified as grid cells (i.e. whose Gridness exceeds the 95^th^ percentile of the distribution of Gridness in their shuffled populations). For all types of Gridness measure, the false positive rate is usually greater than the 5% rate one would expect, confirming our suspicion that current standard methods are susceptible to false positives when applied to populations of cells with spatially inhomogeneous (but irregular) firing patterns. False positive rates are on the whole lower when using field-shuffled rather than spike-shuffled significance thresholds, and diverge less from the expected 5% false-positive rate; thus, indicating a partial solution to the problem.

**Figure 3.**
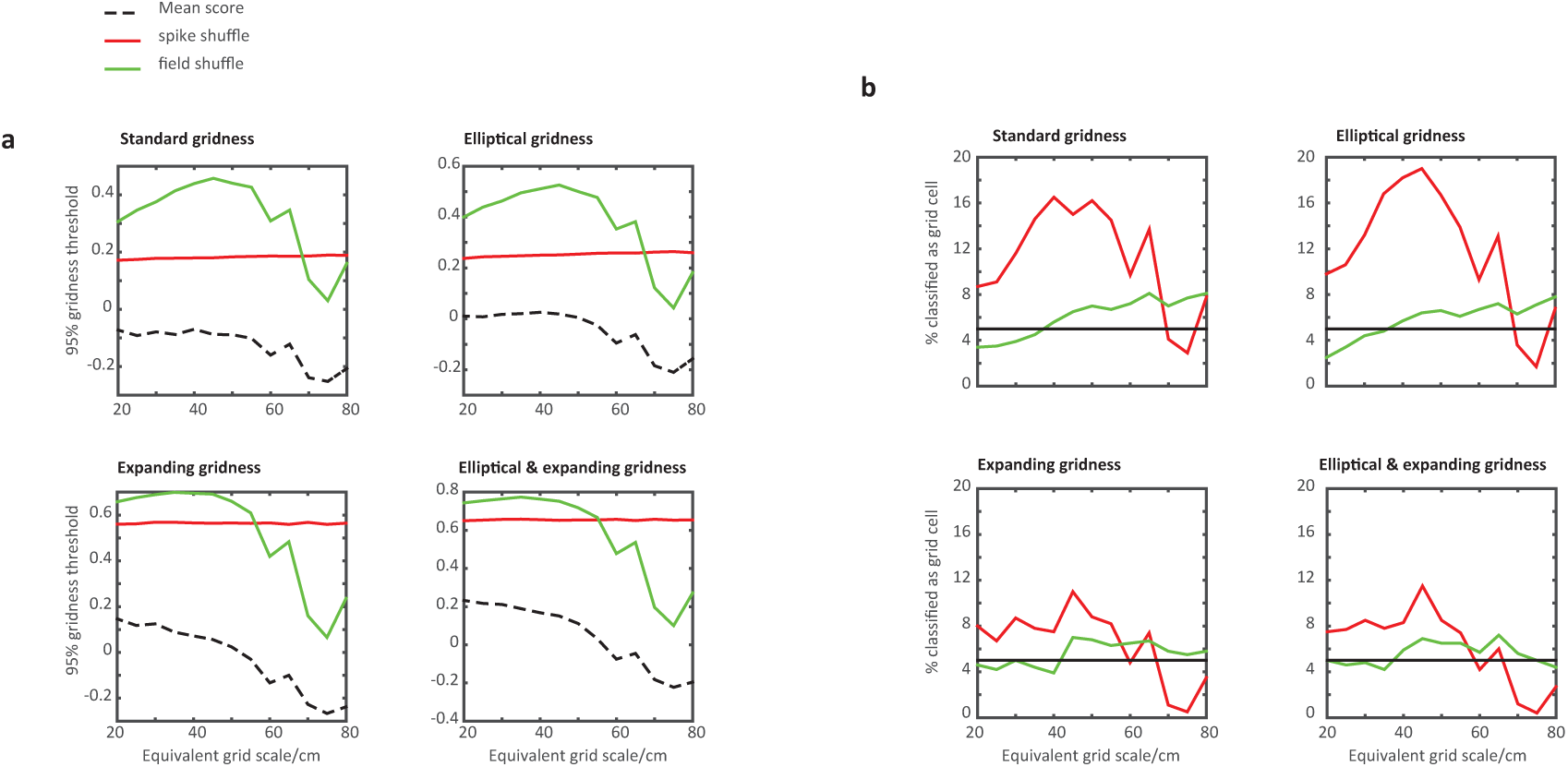
Gridness thresholds and corresponding false positive rates for classifying simulated inhomogeneous irregular firing patterns as “grid cells”. a) 95^th^ percentile Gridness thresholds derived from shuffles of synthetic spatially aperiodic cells under four methods of assessing Gridness. Red, threshold derived from spike shuffling, green from field shuffling, black dotted line is the mean Gridness obtained for each scale in the simulated data. b) Proportion of ‘grid cells’ identified from the irregular simulated firing patterns using the thresholds in (a). When using spike shuffling to establish the null distribution, false positive rates reach ~20% of the elliptical Gridness measure and ~16% for the standard Gridness measure. When using field shuffling, false-positive rates stay much closer to 5% and are generally below 8%.

The simulations show a tendency for the absolute value of the 95^th^ percentile threshold obtained from spike shuffling to be unaffected by the scale of the simulated spatial pattern. Conversely, the threshold obtained for field-shuffles does markedly depend upon the scale, and hence the number of fields within the pattern. In turn, the mean Gridness observed in the synthetic cells also shows a similar dependence on scale; being on the whole lower at larger scales.

The false-positive rate under the spike-shuffle method has a complicated relationship with the size of the simulated pattern; greatly exceeding 5% in many situations. In general, it tends to be highest for spatial patterns with a number of fields equivalent to grid cells with a spatial scale of 40-50cm – having 7.2 and 6.3 fields respectively, on average. Possibly the presence of this number of patches maximises the chance of high “Gridness” despite being irregularly located. The false positive rate under the spike-shuffle method also tends to be slightly lower for both the expanding Gridness based measures, being highest for elliptical Gridness and standard Gridness measures. In contrast to the spike-shuffled method the field-shuffle method exhibits lower false-positive rates at most of the scales tested (Figure 3).

### The Gridness of data from rat medial Entorhinal Cortex

We applied these methods to a sample of single units recorded from the medial Entorhinal Cortex (mEC) of foraging rats. Tetrodes were advanced between recording sessions, ensuring that candidate cells were recorded only once. Similarly, all putative neurons identifiable from their waveforms were manually cluster cut (n=704). Note that these implants were aimed at recording grid cells, and recording terminated when no more grid cells were found, creating a higher proportion of grid cells than might be found in an unbiased sample of cell.

The proportions of active cells that are classified as grid cells in the entire dataset ranges between 22.4% to 24.3% when using a 95% spike shuffled threshold, and between 19.0% to 21.7% with a 95% field shuffled threshold (Figure 4), depending on the exact measure of Gridness used.

**Figure 4.**
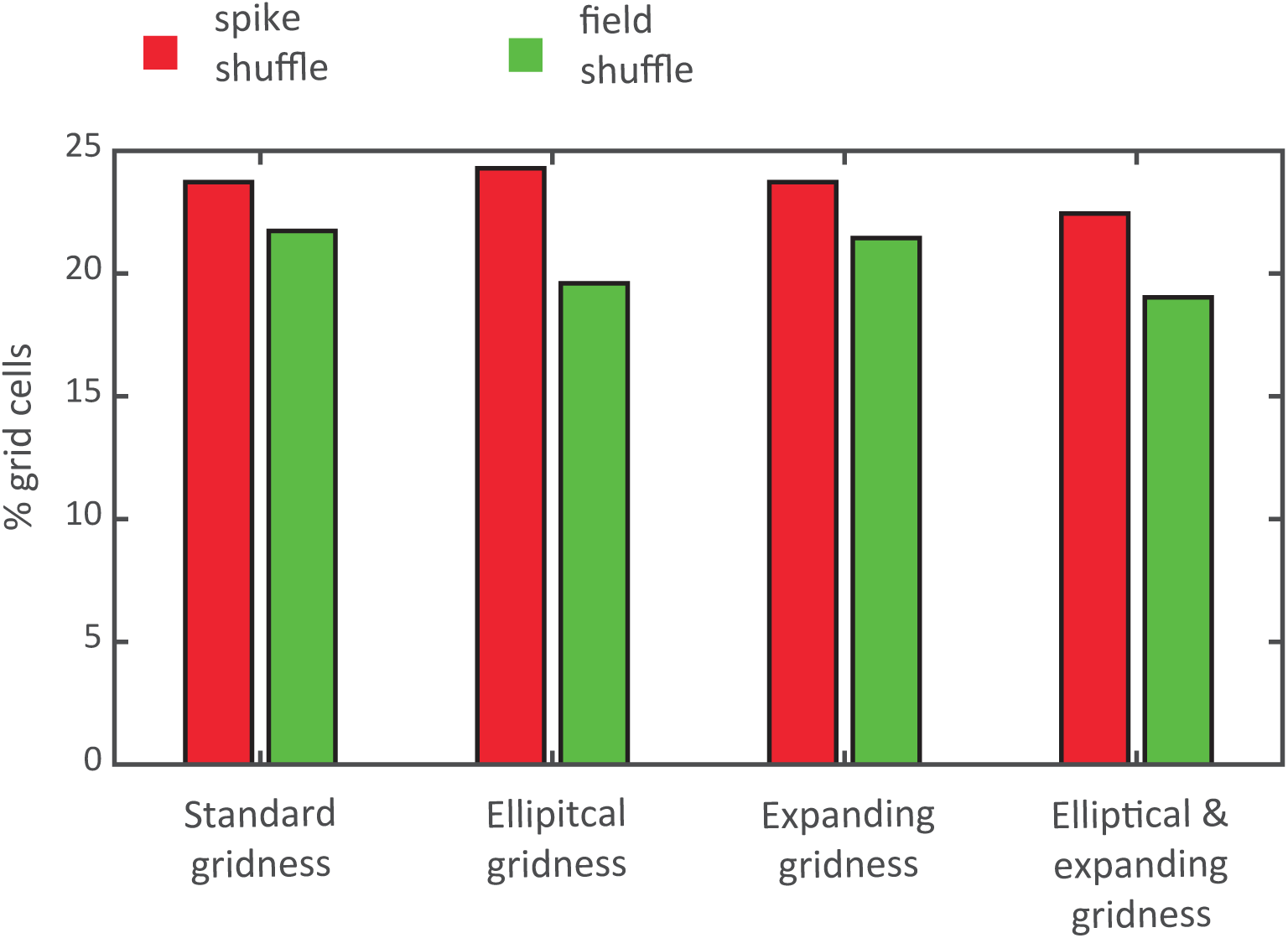
Proportion of grid cells found using the 95^th^ percentile of spike and field shuffled distributions in single unit recordings made from tetrodes located in rat mEC. Red bars indicated the proportion of cells identified on the basis of a spike shuffle and green on the basis of a field shuffle.

## Discussion

Standard methods for detecting grid cells in neural firing data from freely behaving animals involve the use of a statistical threshold based on spike-shuffled data representing the null hypothesis (absence of grid cells). We worried that this method is vulnerable to false-positives caused by the unavoidable spatial inhomogeneity in data from freely-behaving animals. This inhomogeneity arises because movements and perceptual stimuli tend to be inhomogeneously distributed in space during normal behavior, so that neural responses to either could be mistaken for spatial coding (see e.g. Burgess et al. ^15^ regarding the dependencies between directional and locational coding). By contrast, such inhomogeneity is typically not present in spike-shuffled data, reducing its ability to model the null hypothesis for this type of data.

Our simulations demonstrate the potential vulnerability of the standard method for false detection of grid cells when applied to synthetic spatially inhomogeneous (but irregular) firing patterns. We found a surprisingly high false positive rate for a 95% threshold (exceeding 19% in some cases vs. an expected level of 5%). We also showed that using field shuffling rather than spike shuffling to control the false-positive rate in these circumstances has some benefits (the false positive rate is lower, below 8% for the standard Gridness measure). Our simulated false positive rates (Figure 3) may represent a “worse-case” scenario, in that we explicitly chose the spatial inhomogeneity to resemble that of grid cells in the size and number of Gaussian patches, despite their irregular spatial distribution. However, the proportions of cells classified as grid cells with the spike and field shuffle methods did not vary as dramatically in real data from mEC (i.e. a difference of around 2-5%, Figure 4), suggesting that the populations there did not contains such a high proportion of cells with irregular patchy firing patterns.

In conclusion, statistical analysis of “Gridness,” based on the standard spike shuffling method applied to freely behaving animals, are vulnerable to false positive rates potentially approaching 20%, depending strongly on the scale of spatial pattern and type of Gridness measure. The use of spatial field shuffling can reduce the false positive rate somewhat (to less than 8%) and shows a less marked dependence on spatial scale. Thus, the use of field shuffling can be a partial solution to the problem. However, the variability in the Gridness scores for aperiodic spatial patterns of different scales indicates the necessity to generate null distributions on a per cell basis, rather than generating a single threshold for an entire population. In addition, independent corroboration of genuine Gridness is advisable where detected proportions of grid cells approach the potential false-positive rate. Two simple and commonly employed practises are available: to explicitly present all firing patterns, so that the regularity of the pattern can be observed by eye (see Figures 1-2); and to demonstrate that similarly regular patterns generated from the same cell are observable in a second spatial setting and so not a simple confound of the inhomogeneous distribution of a behavioural variable in one setting.

## Acknowledgments

This work was funded by the Wellcome Trust, Royal Society, European Research Council, and Horizon 2020. The authors thank Shailendra Rathore for useful discussion and pilot analyses.

